# Dynamic cerebral autoregulation of endurance-trained men following 6 weeks of high-intensity interval training to exhaustion

**DOI:** 10.1101/605667

**Authors:** Audrey Drapeau, Lawrence Labrecque, Sarah Imhoff, Myriam Paquette, Olivier Le Blanc, Simon Malenfant, Patrice Brassard

## Abstract

Elevated cardiorespiratory fitness (CRF) is associated with reduced dynamic cerebral autoregulation (dCA), but the impact of exercise training *per se* on dCA remains equivocal. In addition, resting cerebral blood flow (CBF) and dCA after high-intensity interval training (HIIT) in individuals with already high CRF is unknown. We examined to what extent 6 weeks of HIIT affect resting CBF and dCA in cardiorespiratory fit men and explored if potential changes are intensity-dependent. Endurance-trained men were assigned to group HIIT_85_ (85% of maximal aerobic power, 1 to 7 min effort bouts, n = 8) and HIIT_115_ (115% of maximal aerobic power, 30 s to 1 min effort bouts, n = 9). Training sessions were completed until exhaustion 3 times/week over 6 weeks. Mean arterial pressure (MAP) and middle cerebral artery mean blood velocity (MCAv_mean_) were measured continuously at rest and during repeated squat-stands (0.05 and 0.10 Hz). Transfer function analysis (TFA) was used to characterize dCA on driven blood pressure oscillations during repeated squat-stands. Neither training nor intensity had an effect on resting MAP and MCAv_mean_ (both *P* > 0.05). TFA phase during 0.10 Hz squat-stands decreased after HIIT irrespective of intensity (HIIT_85_: 0.77 ± 0.22 vs. 0.67 ± 0.18 radians; HIIT_115_: pre: 0.62 ± 0.19 vs. post: 0.59 ± 0.13 radians, time effect *P* = 0.048). These results suggest that HIIT over 6 weeks have no apparent benefits on resting CBF, but a subtle attenuation in dCA is seen posttraining irrespective of intensity training in endurance-trained men.

**NEW & NOTEWORTHY:** The novel findings of this study are that 6 weeks of submaximal and supramaximal high-intensity interval exercise to exhaustion reduce dynamic cerebral autoregulation irrespective of training intensity in endurance-trained men. However, these HIIT protocols do not influence resting cerebral blood flow in these individuals. The results indicate the cerebrovasculature of endurance-trained men has an attenuated ability to react to large and rapid changes in blood pressure following HIIT.

## INTRODUCTION

Established evidence show that cardiorespiratory fitness (CRF) is more cardioprotective when compared to overall physical activity levels (19). CRF-related benefits are not confided to the cardiovascular function and rather extend to the cerebrovascular system (23, 36). For instance, aerobic exercise training alters favorably cerebrovascular health in varying clinical conditions ranging from chronic obstructive pulmonary disease (21), cognitive impairments (6), stroke (14) and following cancer (27). Evidence show that lifelong aerobic training individuals with elevated CRF have higher resting intracranial blood velocity in the anterior circulation, (as indexed by transcranial Doppler sonography of mean blood velocity in middle cerebral artery (MCAv_mean_) (3, 7), and in the posterior circulation (as indexed by arterial spin labelling in posterior cingular cortex/precuneus) (37) and higher extracranial blood flow (as indexed by carotid Doppler) (9) than their sedentary counterparts. In inactive individuals, short-term aerobic training (12 weeks) longitudinally elevates CRF and cerebrovascular reactivity to carbon dioxide, whereas it induces equivocal MCAv_mean_ responses at rest (26). However, whether exercise training can elevate resting CBF in individuals with high CRF remains to be determined.

At rest or in response to challenges such as exercise, a myriad of mechanisms continuously interacts to maintain adequate CBF (4, 35, 40). Among these regulatory mechanisms, dynamic cerebral autoregulation (dCA), i.e. the ability of cerebral vessels to respond to rapid changes in blood pressure (BP), reacts rapidly even before baroreceptor reflex to modulate cerebrovascular resistance and minimize deviations in CBF (1). However, the influence of CRF on dCA remains equivocal. Indeed, while some investigators reported no noticeable differences in dCA between endurance-trained individuals (13) or Masters athletes (2) with untrained individuals, our research group (17) and others (22) observed diminished dCA with elevated CRF. Nonetheless, training-induced effects on dCA in cardiorespiratory fit individuals remain unexplored while dCA seems not to be influenced by aerobic exercise training in older healthy sedentary participants and chronic obstructive pulmonary disease patients (20).

In endurance-trained individuals with already elevated CRF, one option to optimize endurance performance or related physiological adaptations is through the addition of high-intensity interval training (HIIT) (18). HIIT is defined as short bursts of exercise ≥ 80% of maximal heart rate (HR) alternating with periods of recovery or light exercise (38). Requiring half the accumulated time at target intensity, metabolic and cardiovascular health are improved equally and often superiorly with HIIT compared to traditional moderate-intensity continuous training (MICT) (8, 23, 24, 31). Different HIIT protocols are used by athletes, such as submaximal training below maximal oxygen consumption (VO_2max_), training at VO_2max_ and supramaximal training (power output above VO_2max_). Our research group compared two of these HIIT protocols, and recently reported that endurance-trained men performing 6 weeks of submaximal (85% maximal aerobic power) and supramaximal (115% maximal aerobic power) HIIT to exhaustion improve significantly VO_2max_ and anaerobic power irrespective of training intensity (29). However, the knowledge about concurrent cerebrovascular adaptations specific to HIIT conducted at different training intensities remains limited in healthy humans (23).

Therefore, the aim of this study was to examine the influence of submaximal and supramaximal HIIT on resting CBF and dCA in endurance-trained men. We hypothesized that resting CBF would be increased, while dCA would be diminished following HIIT, and the extend of these changes would be intensity-independent.

## MATERIALS AND METHODS

### Ethics and informed consent

The *Comité d’éthique de la recherche de l’IUCPQ-Université Laval* (CER: 20869) approved the study according to the principles established in the Declaration of Helsinki (except for registration in a database). Informed consent was obtained by all participants prior to the investigation.

### Participants

We recruited nineteen endurance-trained men with a history of 5 to 12 h/week for at least 2 years. All participants were free from any diagnosed medical conditions. A variety of endurance sports were undertaken by the participants including road cycling (*n* = 9), triathlon (*n* = 7), mountain biking (*n* = 2) and cross-country skiing (*n* = 1) (29).

### Experimental protocol

This study was part of a larger study examining the influence of submaximal and supramaximal training on determinants of endurance performance (29). However, the current question was determined *a priori* and was prospectively studied as a separate question. For the purpose of our study, we analyzed and compared pre and post-values collected on three visits: 1) anthropometrics, resting systemic and cerebral hemodynamic measurements and the evaluation of dCA 2) incremental cycling test for determination of VO_2max_, and 3) maximal aerobic power evaluation for prescription of training session intensity. Prior to testing, all participants were asked to refrain from consuming alcohol and caffeine for 24 h and to avoid exercise training for at least 12 h. The data were collected in the same order for all participants and the visits were separated by at least 48 h. After being matched according to their age and pre-training VO_2max_, they were randomly assigned to two different intensity training groups; 85% of maximal aerobic power (HIIT_85_) and 115% of maximal aerobic power (HIIT_115_). The post-training testing sessions were repeated 48-96 h following the end of the 6-week training program.

### Training interventions

Over a period of 6 weeks, training consisted in three HIIT sessions per week with 48-72 h between sessions. On the remaining days, participants were allowed to maintain a similar low and/or moderate-intensity volume that they were typically performing prior to the study. Other than the ones already included in the study protocol, HIIT was strictly prohibited.

Specifically, the HIIT_85_ group performed repeated 1-to 7-min efforts bouts. The intensity was set to 85% of maximal aerobic power. The other experimental group, HIIT_115_, repeated 30-s to 1-min effort bouts at 115% maximal aerobic power. Both groups interspaced their effort bout with active recovery (150 W or 50% of maximal aerobic power if maximal aerobic power < 300 W). To avoid routine monotony and to keep the focus on exercise intensity rather than duration, both groups alternated exercise bout duration from one session to another throughout the 6-week period (29). The specificity of our protocol was that both groups had to perform each HIIT session until exhaustion, defined as the inability to complete an effort bout, in order to match the two intensity training protocols for total effort. As already reported, the total training volume was different between groups being 47 % less in HIIT_115_ group compared to HIIT_85_ (19.3 ± 4.7 vs 36.6 ± 14.4 min/session; *P* = 0.005) (29). For further details on the training interventions, refer to (29).

### Measurements

#### Systemic hemodynamics

A 5-lead ECG was used to measure HR. BP was measured beat-to-beat by the volume-clamp method using a finger cuff (Nexfin, Edwards Lifesciences, Ontario, Canada). For uniformity, the cuff was always placed on the right middle finger. BP was corrected by referencing the cuff to the level of the heart using a height correcting unit. The integration of the pressure curve divided by the duration of the cardiac cycle allowed to calculate mean arterial pressure. The dynamic relationship between the BP and cerebral blood velocity is indexed reliably by the volume-clamp method which correlates the dynamic changes in beat-to-beat BP with the intra-arterial BP recordings (28, 32).

#### Middle cerebral artery blood velocity

A transcranial Doppler ultrasound was used at a frequency of 2-MHz pulsed to monitor MCAv_mean_ (Doppler Box; Compumedics DWL USA, Inc. San Juan Capistrano, CA). A standardized procedure (39) was repeated for every participant to localize and identify the left MCA. After the optimal signal was attained, signal depth, gain and power were recorded for post-training evaluations. The probe was fixed to a head set that was placed over a custom-made headband. An adhesive conductive ultrasonic gel (Tensive, Parker Laboratory, Fairfield, NY, USA) was used to ensure a stable position and angle of the probe throughout testing.

#### End-tidal partial pressure of carbon dioxide

A breath-by-breath gas analyzer (Breezesuite, MedGraphics Corp., MN) was used to measure end-tidal partial pressure of carbon dioxide (P_ET_CO_2_) during supine rest baseline and squat-stand maneuvers. Before each evaluation, the analyzer was calibrated to known gas concentrations following manufacturer instructions.

#### Data acquisition

An analog-to-digital converter (Power-lab 16/30 ML880; ADInstruments, Colorado Springs, CO, USA) converted and stored at 1kHz all signals. Subsequent analysis was performed with a free version of a commercially software (LabChart version 8.1.8; ADInstruments).

### Visit 1

#### Anthropometric measurements and resting hemodynamics

Upon arrival, each participant was measured and weighed. Resting hemodynamic measurements included MAP (volume-clamp method using a finger cuff), HR (electrocardiogram), and MCAv_mean_ (transcranial Doppler ultrasound), which were continuously monitored on a beat-by-beat basis in a supine position after 10 min of rest. Cerebrovascular conductance index (CVCi; MCAv_mean_/MAP) and its reciprocal, resistance (CVRi; MAP/MCAv_mean_) was then calculated. Baseline data were averaged over the last 3 min of the resting period, except recordings of P_ET_CO_2_ which were averaged over the last 2 min of the resting period.

#### Assessment of dynamic cerebral autoregulation

dCA was characterized by forcing MAP oscillations using repeated squat-stand maneuvers. It has been shown to be the best reproducible technique to be used and to elicit high interpretable linearity association between MAP and MCAv signals (34). A minimum of 10 min of standing rest was required before performing the squat-stand maneuvers to ensure all cardiovascular variables had returned to baseline. From the standing position, the participants repeatedly squatted down until the back of their legs attained a ~ 90 degrees angle. To achieve a specific frequency of forced oscillations, squat and standing positions were sustained in alternation for a specific time period. In order to achieve the right pace, instructions were given, and participants were asked to practice 2 or 3 squats. Over a period of 5 min, squat-stand maneuvers were repeated at a frequency of 0.05 Hz (10 s squat, 10 s standing) and 0.10 Hz (5 s squat, 5 s standing) (16, 17, 34). Those frequencies were chosen because it has been shown that under the buffering range of 0.20 Hz, large oscillations in MAP are extensively buffered by the cerebral vessels (41). Because it optimizes the signal-to-noise ratio, the squat-stand maneuvers make the interpretations reliable to study cerebrovascular reactivity to BP through a physiologically relevant MAP stimulus. Each participant executed the squat-stand maneuvers at both frequencies (0.05 and 0.10 Hz) in a randomly fashion. A 5-min standing recovery period separated each sequence to assure cardiovascular variables returned to baseline. The breathing instructions to the participants included normal breathing and avoiding Valsalva maneuvers. During this evaluation, MAP, HR, MCAv_mean_, and P_ET_CO_2_ were recorded in a continuous manner. TFA metrics characterizing the linear dynamic relationship between MAP and MCAv is described in more details in the following section. To evaluate whether squats induced changes in P_ET_CO_2_, an averaged P_ET_CO_2_ of the first and last five breaths of each maneuver (0.05 and 0.10 Hz) were calculated (16).

#### Assessment of the dynamic relationship between MAP and MCAv

The recommendations of the Cerebral Autoregulation Research Network (CARNet) (11) were followed when analyzing data using the commercially available software Ensemble (Version 1.0.0.14, Elucimed, Wellington, New Zealand). The spectral analysis of beat-to-beat MAP and MCAv signals was interpolated and re-sampled at 4 Hz. The Welch algorithm was used to do TFA. The analysis required a 5-min recording subdivided into five windows. The successive windows overlapped by 50%. Prior to discrete Fourier transform analysis, data within each subdivision were detrended linearly and passed through a Hanning window.

For TFA, the cross-spectrum between MAP and MCAv was determined and divided by the MAP auto-spectrum to derive the transfer function coherence (fraction of the MAP which is linearly related to MCAv), absolute gain (cm/sec/mmHg) (amplitude of MCAv change for a given oscillation in MAP), normalized gain (%/mmHg), and phase (radians) (difference of the timing of the MAP and MCAv waveforms).

Specific point of estimate of driven frequency (0.05 and 0.10 Hz) were chosen to sample TFA values. They were selected in the very low (0.02-0.07 Hz) and low (0.07-0.20 Hz) frequency ranges where dCA is believed to be the most operant (34). A threshold of 0.50 of coherence was necessary to ensure robustness of the phase and gain subsequent analysis (41). When coherence exceeded the threshold, phase wrap-around was not present.

### Visit 2

#### Maximal oxygen consumption (VO_2max_)

VO_2max_ was determined during a progressive ramp exercise test executed on an electromagnetically braked upright cycle ergometer (Corival, Lode, the Netherlands) (29). Briefly, the exercise protocol included 3 min of resting period followed by a warm-up of 1 min of unloaded cycling, then by an individualized incremental ramp protocol to volitional exhaustion. The highest 30 s averaged VO_2_, concurrent with a respiratory exchange ratio ≥ 1.15 was used to define VO_2max_ (29).

### Visit 3

#### Maximal aerobic power

Maximal aerobic power was measured for the determination of training intensities (85 and 115% maximal aerobic power) as previously described (29).

#### Statistical analysis

The Shapiro-Wilk test was used to confirm normal distribution of data. Data were analyzed by a two-way repeated-measures analysis of variance when normal distribution was confirmed otherwise data were log transformed before analysis. If data were missing, they were analyzed by fitting a mixed effects model. Following detection of an interaction effect (Time x Intensity), differences were identified using within groups paired and between groups independent samples t-tests, with Bonferroni correction. Statistical significance was established *a priori* at *P* < 0.05 for all tests. All values are expressed as mean ± standard deviation. Statistical analyses were performed using a commercially available software (Prism for macOS, version 8, GraphPad software, CA, USA).

## RESULTS

### Participant compliance

A total of 17 out of 19 participants completed the study; HIIT_85_: 26 ± 6 years, 1.77 ± 0.08 m, 72.1 ± 12 kg, *n* = 8 and HIIT_115_: 28 ± 6 y., 1.77 ± 0.09 m, 73.1 ± 7.5 kg, *n* = 9. We excluded two participants from our final analysis; one participant in HIIT_85_ was ill and absent > 3 training sessions and the other participant in HIIT_115_ complained from excessive fatigue during training regime. Excessive noise/artifact in the recordings of two participants (HIIT_85_, *n* = 1; HIIT_115_, *n* = 1), did not allow for TFA measurements, therefore a total of 15 participants were included in the dCA analysis. Of note, we excluded pretraining values of MCAv, CVCi and CVRi of one participant due velocity that did not identify with certainty the MCAv pattern as compared to reference values (39). Also, due to technical reasons (malfunction of the gas exchange analyzer), P_ET_CO_2_ recordings of only 11 participants were completed during baseline rest, 12 participants during pre-training squat-stands and 6 participants post-training.

### Anthropometric measurements and resting hemodynamics

As previously reported by our research group, six weeks of submaximal and supramaximal HIIT did not modify body mass (intensity effect *P* = 0.84; time effect *P* = 0.16; interaction effect *P* = 0.85). However, HIIT protocols significantly increased VO_2max_ in both groups after 6 weeks, demonstrating that the selected training program effectively increased CRF irrespective of intensity (HIIT_85_ pre: 56.0 ± 6.0 vs post: 59.3 ± 4.7 mL/kg/min; HIIT_115_ pre: 55.9 ± 4.0 vs post 59.2 ± 1.1 mL/kg/min, time effect *P* = 0.002) (29).

Concurrently, resting heart rate significantly decreased after training irrespective of intensity (by 8% in HIIT_85_ and 5% in HIIT_115_, time effect *P* = 0.02). However, despite an improvement in CRF, neither training nor intensity resulted in any change in MAP (intensity effect *P* = 0.09, time effect *P* = 0.51; interaction effect *P* = 0.44) and MCAv_mean_ (intensity effect *P* = 0.55, time effect *P* = 0.93; interaction effect *P* = 0.40). CVCi and CVRi were neither influenced by training nor intensity (Table 1).

**Table 1:**
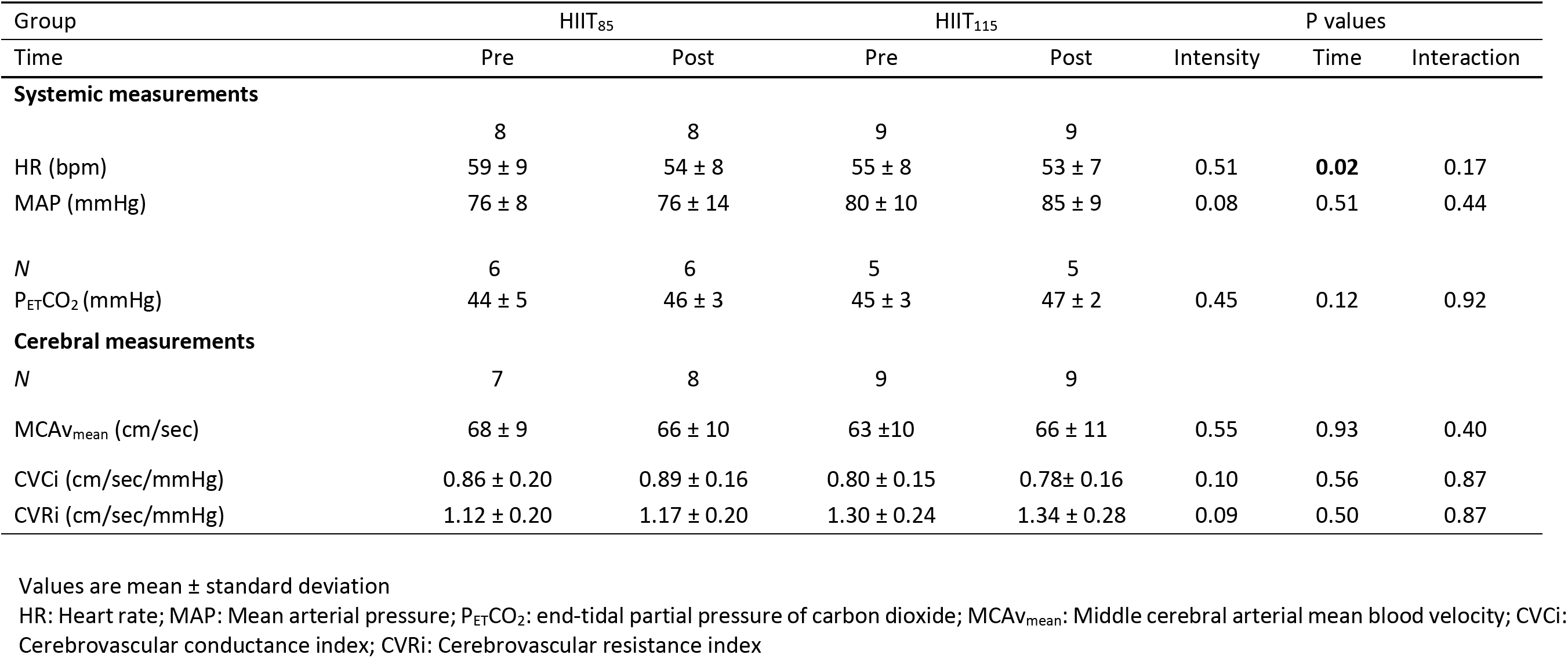
Resting systemic and cerebral hemodynamics at baseline and post-training

| Group | HIIT<sub>85</sub> |  | HIIT<sub>115</sub> |  | P values |  |  |
| --- | --- | --- | --- | --- | --- | --- | --- |
| Time | Pre | Post | Pre | Post | Intensity | Time | Interaction |
| **Systemic measurements** |  |  |  |  |  |  |  |
|  | 8 | 8 | 9 | 9 |  |  |  |
| HR (bpm) | 59 ± 9 | 54 ± 8 | 55 ± 8 | 53 ± 7 | 0.51 | **0.02** | 0.17 |
| MAP (mmHg) | 76 ± 8 | 76 ± 14 | 80 ± 10 | 85 ± 9 | 0.08 | 0.51 | 0.44 |
| *N* | 6 | 6 | 5 | 5 |  |  |  |
| P<sub>ET</sub>CO<sub>2</sub> (mmHg) | 44 ± 5 | 46 ± 3 | 45 ± 3 | 47 ± 2 | 0.45 | 0.12 | 0.92 |
| **Cerebral measurements** |  |  |  |  |  |  |  |
| *N* | 7 | 8 | 9 | 9 |  |  |  |
| MCAv<sub>mean</sub> (cm/sec) | 68 ± 9 | 66 ± 10 | 63 ± 10 | 66 ± 11 | 0.55 | 0.93 | 0.40 |
| CVCi (cm/sec/mmHg) | 0.86 ± 0.20 | 0.89 ± 0.16 | 0.80 ± 0.15 | 0.78 ± 0.16 | 0.10 | 0.56 | 0.87 |
| CVRi (cm/sec/mmHg) | 1.12 ± 0.20 | 1.17 ± 0.20 | 1.30 ± 0.24 | 1.34 ± 0.28 | 0.09 | 0.50 | 0.87 |

Values are mean ± standard deviation
HR: Heart rate; MAP: Mean arterial pressure; P<sub>ET</sub>CO<sub>2</sub>: end-tidal partial pressure of carbon dioxide; MCAv<sub>mean</sub>: Middle cerebral arterial mean blood velocity; CVCi: Cerebrovascular conductance index; CVRi: Cerebrovascular resistance index

### TFA of forced oscillations in MAP and MCAv

The power spectrum densities of MAP and MCAv during repeated squat-stands at 0.05 and 0.10 Hz did not differ between groups or across time (pre- and post-training) (all *P* > 0.05; Table 2). HIIT decreased TFA phase during 0.10 Hz squat-stands irrespective of training intensity (effect of time *P* = 0.048; Figure 1). TFA gain and coherence were unaffected by training or intensity during repeated squat-stands at both frequencies (all *P* > 0.05). There were no significant changes in P_ET_CO_2_ during repeated squat-stands at 0.05 Hz and 0.10 Hz (all *P* > 0.05) and were comparable between groups and time (pre-vs. post-training; Table 2).

**Figure 1.**
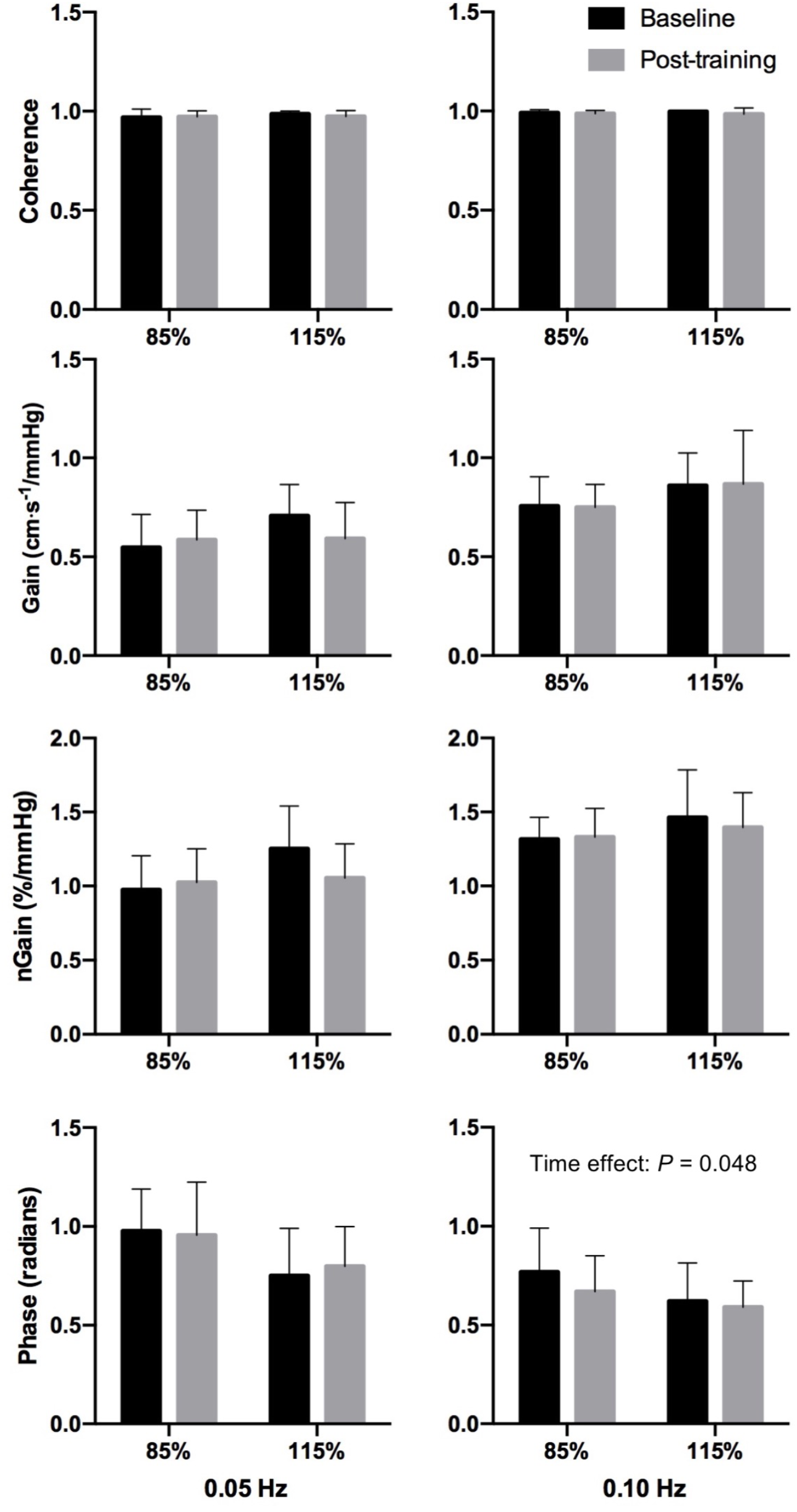
Transfer function analysis of forced oscillation in mean arterial pressure and middle cerebral artery blood velocity. Group averaged coherence, gain, normalized gain (nGain) and phase for 0.05 and 0.10 Hz repeated squat-stands.

**Table 2:** Power spectrum densities and change in end-tidal partial pressure of carbon dioxide during forced oscillations in mean arterial pressure and middle cerebral artery blood velocity during squat-stand maneuvers

| Groupe | HIIT<sub>85</sub> |  | HIIT<sub>115</sub> |  | Valeurs P |  |  |
| --- | --- | --- | --- | --- | --- | --- | --- |
| Time | Pre | Post | Pre | Post | Intensity | Time | Interaction |
| **Squats 0.05 Hz** |  |  |  |  |  |  |  |
| *N* | 8 | 8 | 9 | 9 |  |  |  |
| MAP power (mmHg<sup>2</sup>) | 19424 ± 19255 | 18644 ± 19614 | 10814 ± 7845 | 14325 ± 13938 | 0.41 | 0.65 | 0.48 |
| MCAv power (cm/s<sup>2</sup>) | 9217 ± 13070 | 8936 ± 13572 | 5203 ± 3539 | 6488 ± 9749 | 0.55 | 0.76 | 0.63 |
| *N* | 6 | 3 | 6 | 3 |  |  |  |
| P<sub>ET</sub>CO<sub>2</sub> (Δ mmHg) | -2 ± 3 | -2 ± 4 | 2 ± 1 | 1 ± 2 | 0.26 | 0.16 | 0.31 |
| **Squats 0.10 Hz** |  |  |  |  |  |  |  |
| *N* | 8 | 8 | 9 | 9 |  |  |  |
| MAP power (mmHg<sup>2</sup>) | 16130 ± 13640 | 15895 ± 15144 | 14281 ± 10764 | 14510 ± 12134 | 0.80 | 1.00 | 0.91 |
| MCAv power (cm/s<sup>2</sup>) | 9715 ± 8263 | 8682 ± 8416 | 7966 ± 5059 | 9208 ± 8806 | 0.88 | 0.94 | 0.39 |
| *N* | 6 | 3 | 6 | 3 |  |  |  |
| P<sub>ET</sub>CO<sub>2</sub> (Δ mmHg) | 1 ± 4 | 1 ± 2 | 3 ± 2 | 3 ± 4 | 0.18 | 0.99 | 0.77 |

Values are mean ± standard deviation
HR: Heart rate; MAP: Mean arterial pressure; MCAv: Middle cerebral arteria; blood velocity; P<sub>ET</sub>CO<sub>2</sub>: end-tidal partial pressure of carbon dioxide

## DISCUSSION

To our knowledge, our study is the first to assess the longitudinal effects of submaximal and supramaximal HIIT to exhaustion on resting cerebral hemodynamics and dCA in endurance-trained men. The main findings of this study are threefold: 1) resting CBF remained unchanged after 6 weeks of HIIT; 2) TFA phase during 0.10 Hz repeated squat-stand maneuvers was reduced following training irrespective of intensity; 3) training intensity did not have any effect on our main outcome variables.

### Effect of HIIT on resting cerebral blood flow

We found resting MCAv_mean_ remained unchanged following 6 weeks of submaximal or supramaximal HIIT to exhaustion. Our results differ from cross-sectionnal studies assessing life-long training effect that reported elevated resting MCAv_mean_ (3, 7) and regional CBF in the posterior cingular cortex (37) in elderly trained participants. Longitudinally, short-term aerobic exercise of varied nature (continuous or interval training), intensity and duration [12 weeks (6, 26) and 8 weeks (20)] has been associated with improved CRF in healthy individuals. In association with this training-induced CRF improvement, some investigators reported a small but significant increase in MCAv_mean_ following a 12-week program (26), whereas others did not observe any change in MCAv_mean_ (20) or whole-brain CBF (6) after respectively 8 and 12 weeks of training.

Nevertheless, the effect of exercise training using solely HIIT to exhaustion had not yet been documented in young endurance-trained men with already high CRF. In light of our results, it is possible that training duration might explain discrepancies. Indeed, 6 weeks of exercise training may not be a long enough stimulus to instigate noteworthy and clear beneficial adaptations (i.e. elevation in resting CBF) in young individuals, although HIIT is assumed to provide enough stimulus for cerebrovascular adaptation to occur (23). Considering that it has been argued that the brain vasculature needs to be challenged in order to witness adaptive changes in cerebral hemodynamics (10), our results do not rule out a potential influence of HIIT on CBF changes during aerobic exercise.

### Effect of HIIT on dynamic cerebral autoregulation

We have previously demonstrated cross-sectionally that dCA is impaired in endurance-trained men with elevated CRF (17). In that report, we showed that elevated CRF was associated with increased TFA gain during 0.10 Hz squat-stands compared to sedentary individuals, suggesting a subtle attenuation in dCA. In the present study, we trained a sample of the men included in that cross-sectional study by Labrecque et al. (2017) over 6 weeks with HIIT to exhaustion and investigated whether dCA would be further impaired, and whether these potential changes would be influenced by training intensity. The current findings revealed that irrespective of training intensity, 6 weeks of HIIT did not further impair TFA gain in these participants (unchanged TFA gain post-training *P* > 0.05), but rather decreased TFA phase during repeated squat-stands at 0.10 Hz, irrespective of training intensity. These results suggest that when endurance-trained men with already elevated VO_2max_ further improve CRF with HIIT, they also aggravate their attenuated dCA [TFA gain during 0.10 Hz squat-stands previously known to be higher in these endurance-trained men vs. sedentary controls (17)] through a reduction in TFA phase. The clinical and physiological implications of this subtle attenuation in dCA following HIIT in endurance-trained individuals remain to be elucidated.

To our knowledge, only one other longitudinal study assessed dCA following aerobic exercise training of varied nature and intensity (20). These authors reported no training effect on dCA in healthy older sedentary participants. One possible explanation might be that despite an ~18% increase in CRF, the sedentary older participants (>64 years) studied by Lewis et al. (2019) had significantly lower CRF levels at baseline than our young endurance-trained men (26 to 28 years). These results support that elevated CRF is associated with diminished dCA, whereas dCA is maintained in individuals with lower CRF (20). Further research is required to elucidate whether a threshold exists above which CRF will be associated with an attenuated dCA. In addition, Phillips et al. (2018) have recently reported that 4 weeks of repeated transient hypertension induced by colorectal distension in rats after spinal cord injury lead to impairments of the brain vasculature such as cerebrovascular endothelial dysfunction and profibrotic cerebrovascular stiffness (30). Unfortunately, we did not monitor BP changes during HIIT sessions to exhaustion performed 3 times/week for 6 weeks. We could speculate the subtle attenuation in dCA observed following HIIT in our participants is related to subclinical cerebrovascular impairments induced by the repetitive and rapid surges in BP during each HIIT session performed over 6 weeks.

### Limitations

The findings from our present work should be interpreted in view of limitations. We acknowledge the uniqueness of our exhaustive HIIT training protocols limited the number of endurance-trained participants which resulted in a small sample size. Nonetheless, it allowed us to explore the particularities, feasibility and longitudinal effects of training in this specific population. We relied on transcranial Doppler ultrasonography to measure cerebral blood velocity. This method is only valid if the insonated artery diameter remains constant. It is likely that our study protocol induced negligible physiological range of variation of MAP and P_ET_CO_2_ that could impact on the diameter of the artery (5). The use of transcranial Doppler remains recognized to provide excellent temporal resolution and has been validated as an indirect surrogate marker of CBF (33). Finally, considering that sex (12, 16) and various clinical conditions such as type 2 diabetes and pulmonary arterial hypertension (15, 25) have an influence on dCA, our findings cannot be generalized to healthy women or patients.

## CONCLUSION

Collectively, these findings suggest that 6 weeks of HIIT to exhaustion does not influence resting CBF but is associated with a subtle attenuation in dCA irrespective of training intensity in endurance-trained men.

## GRANTS

Funding of the current project came from the Ministère de l’Éducation, du Loisir et du Sport du Québec and the Fundation of the Institut Universitaire de Cardiologie et Pneumologie de Québec. A scholarship was granted to Myriam Paquette from the Canadian Institute of Health Research. The Société Québécoise d’Hypertension Artérielle supported LL with a doctoral scholarship and SI has her doctoral training supported by the Fonds de recherche du Québec.

## DISCLOSURE

No conflicts of interest, financial or otherwise, are declared by the author(s).

## ACKNOWLEDGMENTS

A special thank you to all the participants that agreed to perform our exhaustive training protocol. Without them, the research could not have been done. We received special assistance from Louis-Charles B. Lacroix and Andrée-Anne Clément, who supervised the training sessions. We are grateful for Cycle Lambert who generously lended us the home trainers Tacx Bushido.

## Authors contribution

P.B. contributed to the original idea of the study; M.P., O.L.B., S.M. and P.B. contributed to data collection; A.D., L.L. contributed to data analyses; A.D., L.L., S.I. contributed to data interpretation; A.D. and P.B. drafted the article. All authors provided approval of the final article.

